# A behavioral logic underlying aggression in an African cichlid fish

**DOI:** 10.1101/2020.07.22.216473

**Authors:** Beau A. Alward, Phillip H. Cathers, Danielle M. Blakkan, Russell D. Fernald

## Abstract

Social rank in a hierarchy determines which individuals have access to important resources such as food, shelter, and mates. In the African cichlid fish *Astatotilapia burtoni*, rank is under social control, such that larger males are more likely than smaller males to be dominant in rank. Although it is well known that the relative size of *A. burtoni* males is critical in controlling social rank, the specific behavioral strategies underlying responses to males of different sizes are not well understood. In this research, our goal was to characterize these responses by performing resident-intruder assays, in which aggressive behaviors were measured in territorial males in response to the introduction of unfamiliar males that differed in relative standard length (SL). We found that the relative SL of intruders played an important role in determining behavioral performance. Resident males exposed to larger (>5% larger in SL) or matched (between 0 and 5% larger or smaller in SL) intruder males performed more lateral displays, a type of non-physical aggression, compared to resident males exposed to smaller (>5% smaller in SL) intruder males. However, physical aggression, such as chases and bites, did not differ as a function of relative SL. Our results suggest that *A. burtoni* males amplify non-physical aggression to settle territorial disputes in response to differences in relative SL that were not previously considered to be behaviorally relevant.

**Highlights:**

- Relative size determines social rank in the African cichlid *Astatotilapia burtoni*
- Resident male *A. burtoni* respond differently to small size differences in intruder males
- Residents perform more non-physical aggression against larger intruders
- Residents do not alter physical aggression as a function of differently sized intruders
- Distinct behavioral strategies are used against different intruders

## Introduction

Intraspecific aggression is widespread among social animals (van Staaden, Searcy, & Hanlon, 2011). Aggressive behavior, either through physical attacks or non-physical signaling, is used to resolve conflicts related to access to resources such as food, shelter, territory, and mates. Extraordinary diversity exists in how different species express aggression and the rules that govern aggressive interactions. However, one rule seems to apply across species: physical or injurious behaviors are considered to be escalatory, occurring primarily in response to conflicts that are difficult to resolve (Holekamp & Strauss, 2016; Maynard Smith & Harper, 1988; van Staaden et al., 2011). The degree of conflict has been formally defined in terms of differences in resource holding potential (RHP). RHP can take the form of different levels of fighting ability as measured by body or weapon size. When large asymmetries in RHP exist among animals, aggressive interactions do not escalate from non-physical to physical; however, when asymmetries in RHP among animals are smaller, aggressive interactions are more likely to escalate, involving more physical and injurious forms of aggression.

Evolution has shaped social dynamics across species to resolve aggressive interactions with as little physical fighting as possible, as this ensures individual and species survival (Holekamp & Strauss, 2016; Maynard Smith & Harper, 1988; van Staaden et al., 2011). This is abundantly clear in social animals that exist in a hierarchy, where rank determines which individuals possess a territory and the behaviors they perform. This is the case for the African cichlid fish *Astatotilapia burtoni*, where males stratify along a dominance hierarchy and exist as either non-dominant or dominant (Fernald, 2012). Dominant males possess a territory which they defend through aggressive interactions and in which they mate with females, while non-dominant males do not perform these behaviors. Dominant males also possess larger testes and brighter body coloration compared to non-dominant males. Social hierarchies in *A. burtoni* remain in flux, however, as non-dominant males constantly survey the environment, searching for a social opportunity to ascend in social rank to dominance. Social opportunity for a non-dominant *A. burtoni* male typically occurs when a larger male is absent from the environment, which a given smaller non-dominant male perceives as an opportunity to ascend to dominant rank. Within minutes of the opportunity, the non-dominant male increases aggressive and reproductive behavior in an attempt to establish a territory. Dominant males who encounter a larger dominant male in their environment will begin to descend in social rank by reducing aggressive and reproductive behavior (Maruska, Becker, Neboori, & Fernald, 2013).

Size-induced social control of social status in *A. burtoni* has been shown in several studies. The reliable occurrence of this phenomenon makes size an excellent tool for controlling social environments in the laboratory, with the goal of generating fish with a given social status and studying the associated physiological underpinnings (for examples, see Alward, Hilliard, York, & Fernald, 2019; Maruska, Becker, Neboori, & Fernald, 2013; Maruska & Fernald, 2010). Although it has been shown repeatedly that size influences social status in male *A. burtoni*, a precise understanding of the relationship between size and behavior has not been established. For instance, while size is something that modifies social decisions in male *A. burtoni*, it is unclear what size difference males actually perceive as different and how they modify their behavior accordingly. Previous work has defined male *A. burtoni* as “matched” in size within a large range of standard length (e.g., 0-10% larger or smaller in standard length (SL=measured from the most anterior portion of the mouth to the most anterior portion of the caudal fin); see Alcazar, Hilliard, Becker, Bernaba, & Fernald, 2014; Desjardins & Fernald, 2010)). However, recent work suggests that very small size differences between male *A. burtoni* can affect social interactions. For example, Alcazar et al (2014) found that males that were 2.1-4.9% larger in SL than their competitor consistently won during a contest, suggesting that size differences previously regarded as “matched” may actually be behaviorally relevant. However, this study was focused on which fish won each contest and not on the specific behavioral strategies underlying responses to differently sized males.

Characterizing the specific behavioral patterns in *A. burtoni* that occur in response to differently sized males may yield insight into the capacity of *A. burtoni* to discern different levels of social opportunities, which would allow for a deeper understanding of the cognitive abilities required to successfully navigate a social hierarchy. In the present study we characterized behavioral responses in male *A. burtoni* as a function of differently sized male competitors during resident-intruder assays, in which a dominant male with a territory (i.e., the resident) was exposed to an unfamiliar, non-dominant male intruder that differed in relative standard length (SL). (illustrated in Fig. 1). The results of this study could shed light on the rules of engagement during social interactions in male *A. burtoni*.

**Figure 1.**
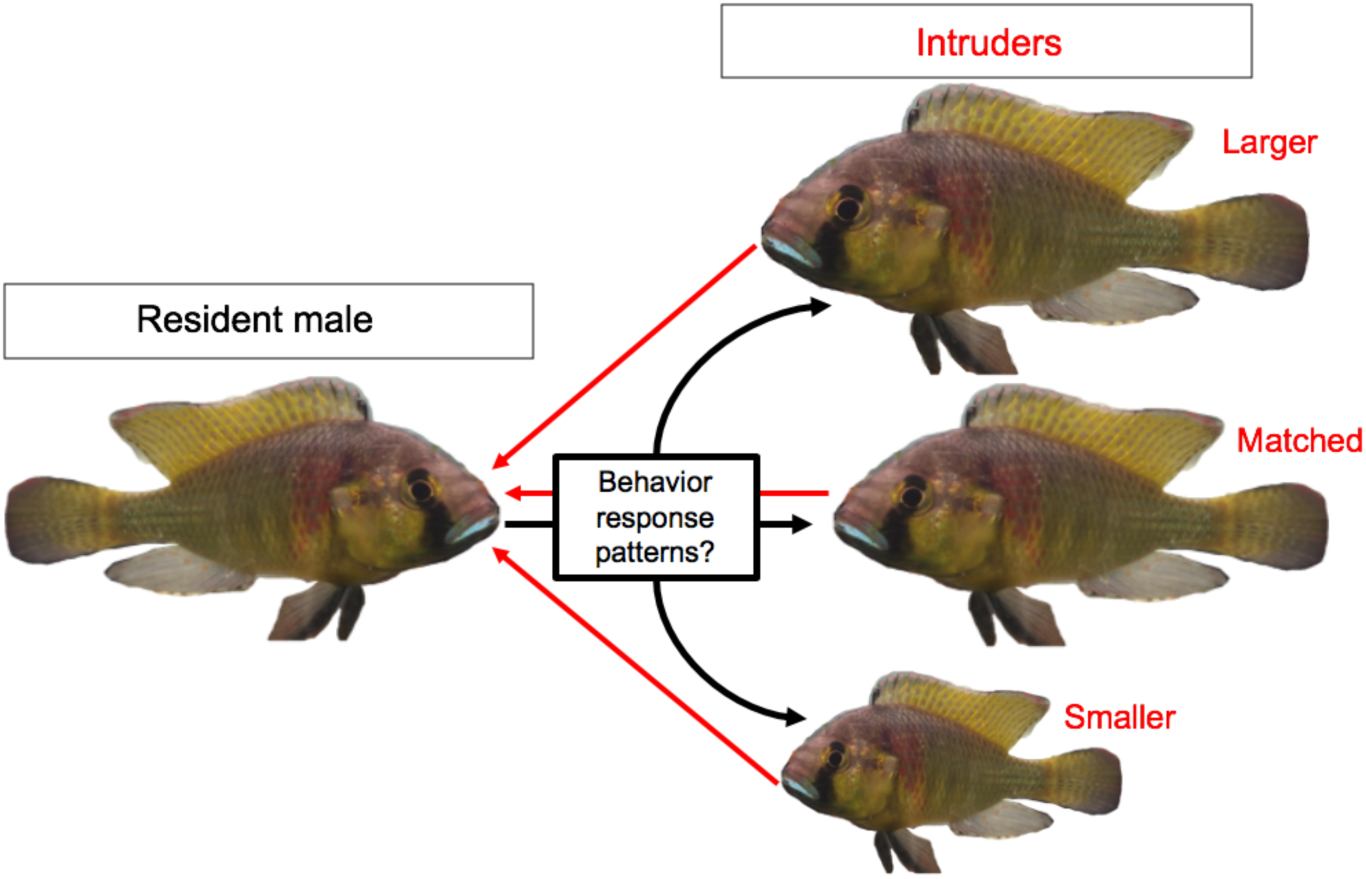
The behavioral patterns in male *Astatotliapia burtoni* underlying responses to differently sized males have not been characterized. Male *A. burtoni* change their social status depending on the social environment. Large males socially suppress smaller males and large males are more likely to be dominant. The specific behaviors males perform in response to differences in relative size, however, have not been determined. We asked during a resident-intruder assay what behavior patterns resident males use when presented with male intruders that were smaller, larger, or matched in size.

## Methods

### Ethical Note

The protocols and procedures used here were approved by the Stanford University Administrative Panel on Laboratory Animal Care (protocol number: APLAC_9882) and followed the ASAB/ABS Guidelines for the use of animals in research. We were able to monitor the behaviours of all fish throughout each day of the study (see below). Throughout the whole assay, each tank was monitored in real time through a Wi-Fi-enabled camcorder remotely connected to a tablet (iPad). Fish in all other tanks were monitored three times daily by visual inspection, to ensure they experienced no physical harm. No fish were physically harmed at any point during the assay.

### Animals used

Fish were bred and used at Stanford University from a colony derived from Lake Tanganyika in accordance with AAALAC standards.

### General approach

To assess behavior as a function of SL, we conducted resident-intruder assays. We took several steps to control for social experience and age of both the resident and intruder, since these factors have been found to influence behavior in *A. burtoni* (Alcazar et al., 2014;

Solomon-Lane & Hofmann, 2019). In these assays, the resident had a dominant social status and an established territory, while the intruder had a non-dominant social status. To control for previous social experience, the resident and intruder were unrelated and had had no visual, physical, or chemical interaction at any point prior to the assay. Another step we took to control for social experience was to socially suppress all males before they were given the social opportunity to ascend to dominance and were provided a territory. We also physically isolated socially ascending fish from other fish, to further control for the role of social experience on behavioral responses to the intruder. All intruders were socially suppressed in a tank with fish that were unrelated to the resident. Finally, all resident-intruder pairs were age-matched, to control for effects of age on behavior (Alcazar et al., 2014).

### Assay set up

#### Social suppression

Two 121-liter social suppression tanks (see Fig. 2a for illustrated example) were each filled with 20 related, small suppressed males, as well as 3 large, unrelated dominant males and 3 females. The two tanks contained broods of the same age from different parents. Fish from the two tanks could not interact visually, physically, or chemically with those in the other tank. Smaller suppressed males were housed in these conditions for at least 45 days before being transferred to a dominance inducing tank (see below).

**Figure 2.**
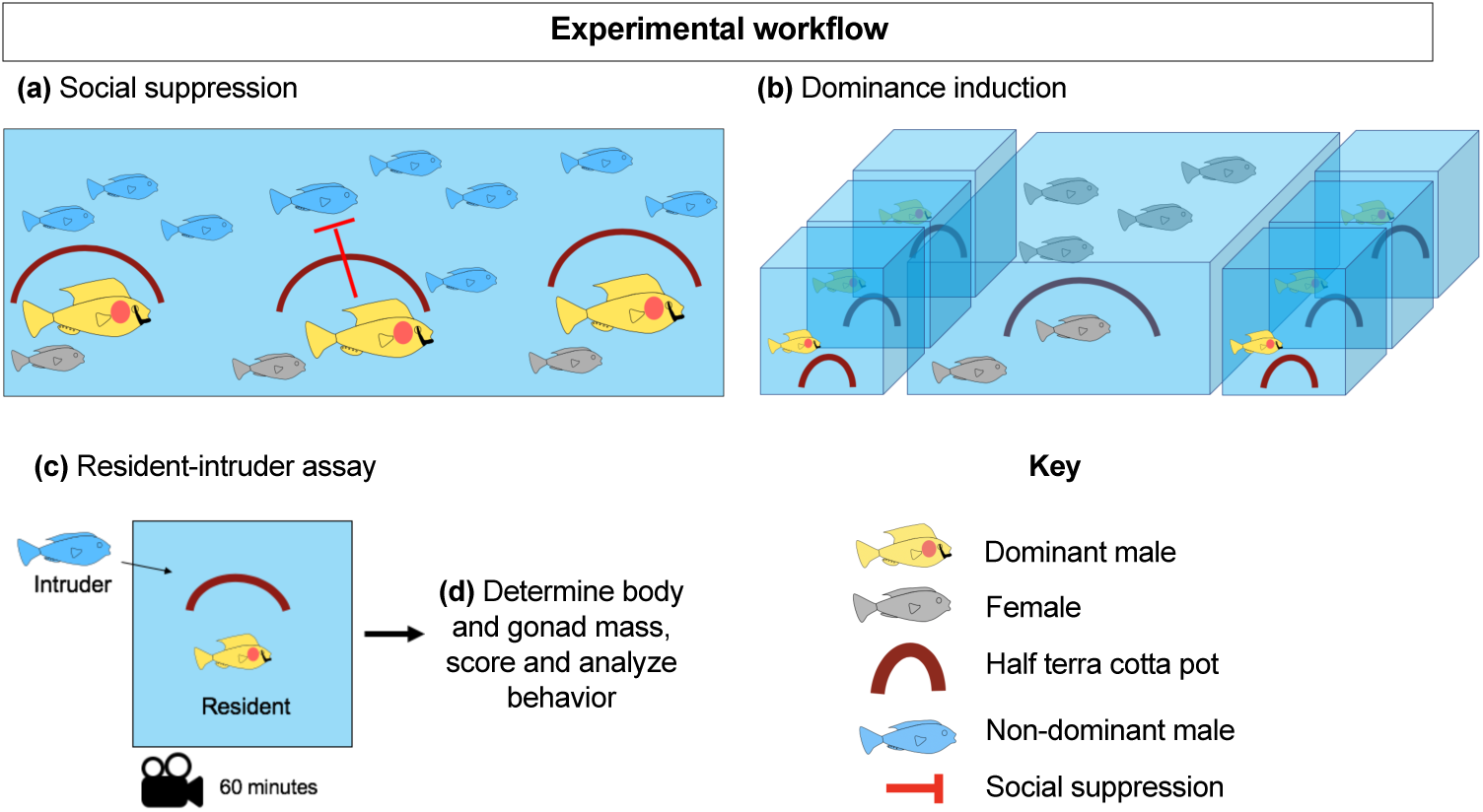
Experimental workflow to probe behavioral strategies in male *Astatotilapia burtoni*. (a) Males were socially suppressed to non-dominant social status using social suppression tanks. (b) After social suppression, males were placed individually into dominance inducing tanks where they possessed and could interact visually with males and females. (c) Once males reach social dominance, they are transferred to a tank where they establish a territory as a resident and then exposed to a male intruder. (d) After the assay, male residents are weighed for body and gonad mass and behavior videos are scored and analyzed.

### Dominance inducing tank setup

Thirty-liter dominance inducing tanks (see Fig. 2b for illustrated example) were set up for the isolation of previously suppressed males to allow for controlled social ascent to dominance. Each tank contained a shelter (half terra cotta pot) and faced a 121-liter tank filled with 10 unrelated females with which the male could interact visually but not physically or chemically. Beside each tank was another isolated male in a dominance-inducing tank with which they could interact visually but not physically or chemically. Males were transferred from the suppression tank and isolated in a dominance-inducing tank for 2-4 weeks before entering the assay tank.

### Resident-intruder assay

A resident-intruder assay tank (see Fig. 2c for illustrated example) consisted of one 30-liter tank, with a half terra cotta pot (Fig. 2c). A male was removed from a dominance-inducing tank, its SL was measured, and immediately placed in an 30-liter tank containing gravel and a half terra cotta pot simulating a spawning site. The male was given 48 hours to acclimate and establish a territory (Alward et al., 2019).

After the 48-hour acclimation period, an intruder male was removed from its social suppression tank, its SL was measured, and then it was introduced to the resident-intruder assay tank. Video recording began as soon as the intruder was introduced. Behavioral interactions were also monitored remotely in real time (iPad). Immediately after observation of the first aggressive behavior by the intruder or resident, recording continued for 60 min more and then the assay was stopped.

### Dissections

Immediately following the completion of the resident-intruder assay, the resident male was removed, weighed, and euthanized via rapid cervical transection (see Fig. 2d). An incision was made anterior to the vent to the caudal fin and the gonads were removed and weighed.

### Scoring behavior

Based on previous work, multiple types of behavior were quantified (Fernald & Hirata, 1977; see Fig. 3 for illustrated examples of behaviors): fleeing from male; physical aggression (chase male and bite male); non-physical aggression (lateral display and flexing); and pot entry, a territorial behavior. Fleeing was defined as a rapid swim retreating from an approaching fish. Chase was defined as a rapid swim directed towards a fish. Biting was defined as the male lunging a short distance towards a fish and biting it on its side, then floating backwards a short distance. Lateral displays were defined as aggressive displays classified as presentations of the side of the body to another fish with erect fins, flared opercula, and trembling of the body. Flexes were defined as presentations of the side of the body with erect fins while the fish was immobile. Pot entry was defined as as any time a male entered the half terra cotta pot. Videos were scored in Scorevideo (Matlab). The results of scoring videos were saved into log files that were subjected to a variety of analyses using custom R software.

**Figure 3.**
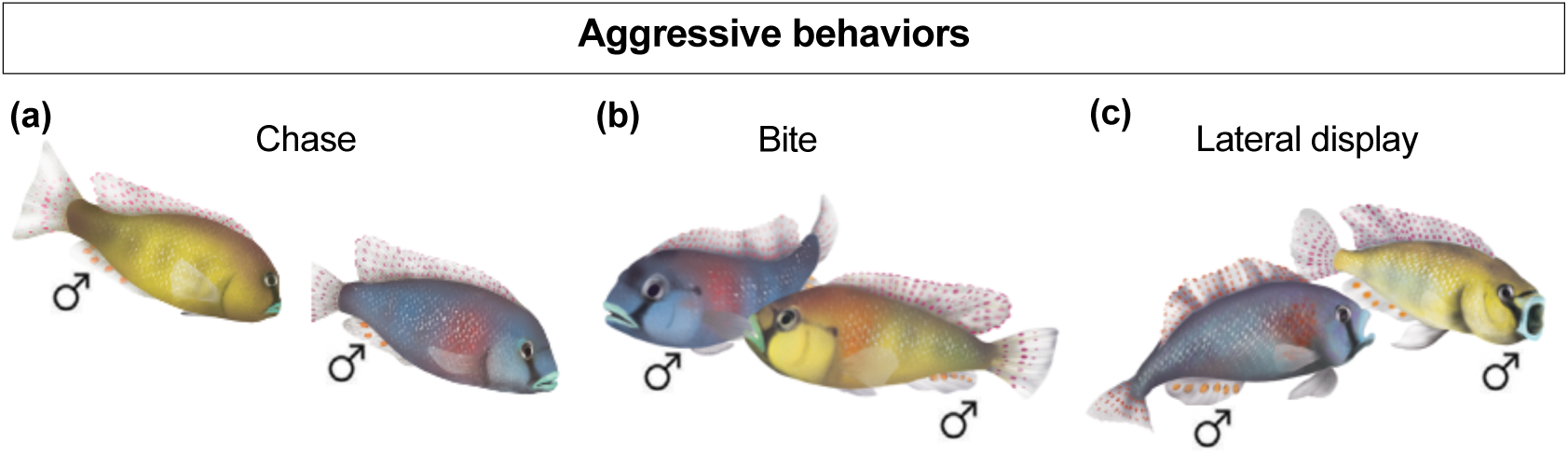
Illustration of aggressive behaviors. We quantified multiple aggressive behaviors performed during resident-intruder assays, including (a) chases, (b) bites, and (c) lateral displays directed towards males.

### Measuring the effects of size differences: group-level and continuous analyses

Previous work in *A. burtoni* and other cichlids suggests that a size difference between 0 and 5% is considered “matched” in size (Alcazar et al., 2014; Reddon et al., 2011; Taborsky, 1984, 1985). Therefore, to explore the behavioral strategies used as a function of relative SL (intruder SL/resident SL), we used three groups for our resident-intruder assays: Smaller, Matched, and Larger. The “Smaller” group contained residents that were exposed to intruders 5% or more larger in SL; the “Matched” group included residents that were within ±5% of the size of the intruder; and the “Larger” group included residents that were exposed to intruders 5% or more smaller in SL. We also assessed the effects of relative SL as a continuous variable on behavior using correlational analyses.

### Statistical analysis

All statistical tests were performed in Prism 8.0. We used Kruskal-Wallis ANOVAs followed by Dunn’s post-hoc tests for comparisons of physiological and behavioral measures across groups. When comparing only two groups, we used Mann-Whitney tests. Raster plots were generated using custom software packages in R (available at https://github.com/FernaldLab). Correlational analyses were conducted using Pearson’s *r*. Effects were considered significant at *p*≤0.05.

## RESULTS

### Qualitative analysis of behavior as a function of relative SL

We first visualized behavioral output for all fish in raster plots (Fig. 4). These plots showed that regardless of relative SL, residents attacked intruders at similar rates. However, fish from both the Matched (Actual SL difference range=intruder 0 to 4% larger than resident) and the Smaller group (Actual SL difference range=intruder 5 to 10% larger than the resident) performed more lateral displays than the Larger group (Actual SL difference range=intruder 5 to 23% smaller than the resident), suggesting that as intruder SL increases relative to the resident, residents perform more non-physical acts of aggression. Finally, most intruders performed zero aggressive behaviors towards the resident (see Fig. 5; results not shown in raster plots because it occurred at such low rates and only in a few fish; see below), indicating that the resident fish all maintained dominance throughout the challenge (or “won”).

**Figure 4.**
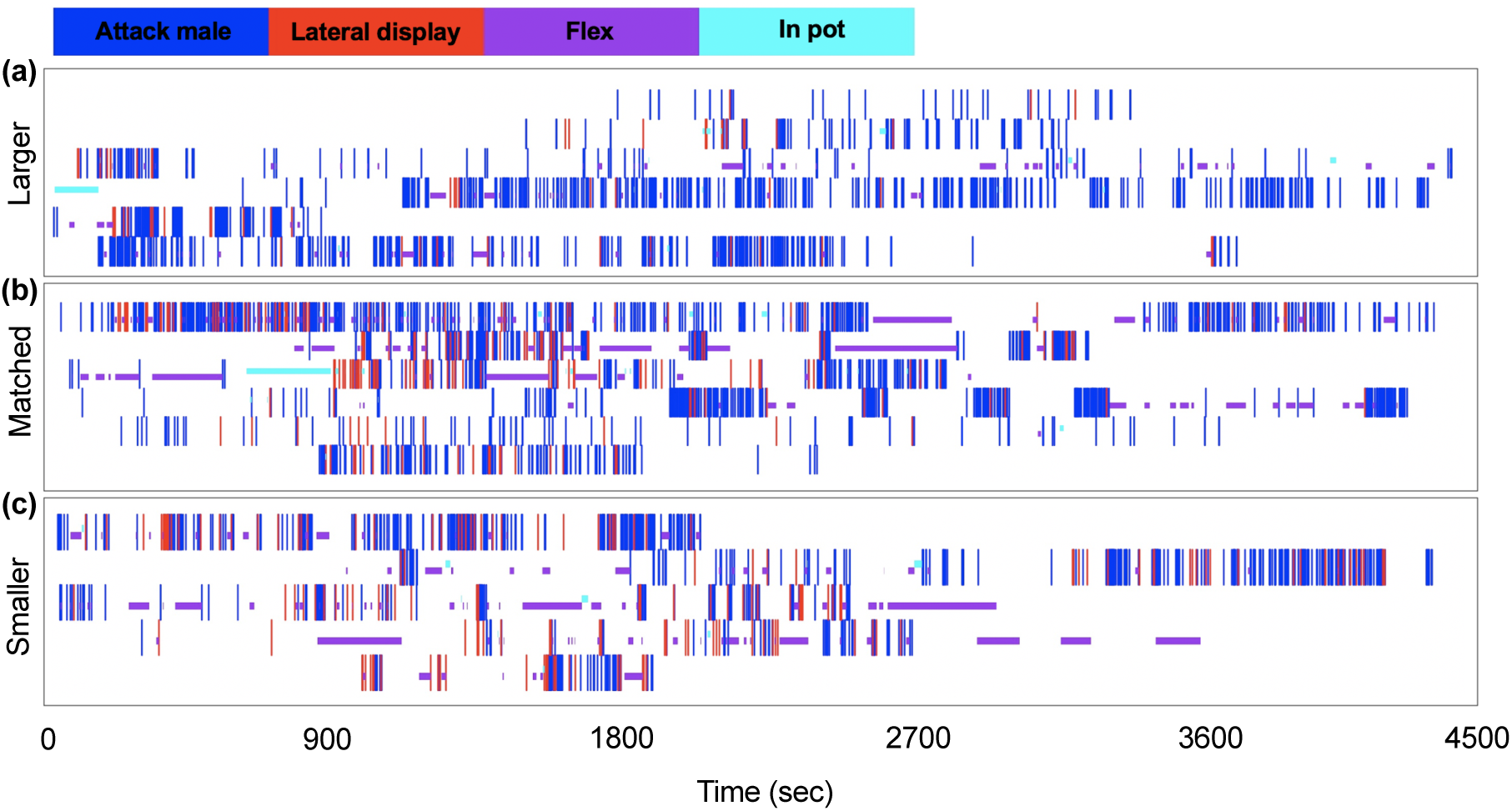
Qualitative visualization of behavior. Raster plots showing behavior from individual fish from each the (a) Larger, (b) Matched, or (c) Smaller group. Each colored line represents a particular type of behavior. The x-axis represents time. “Flex” and “In pot” are represented here are durational behaviors; once the bar denoting either behavior is over the fish has stopped that behavior.

**Figure 5.**
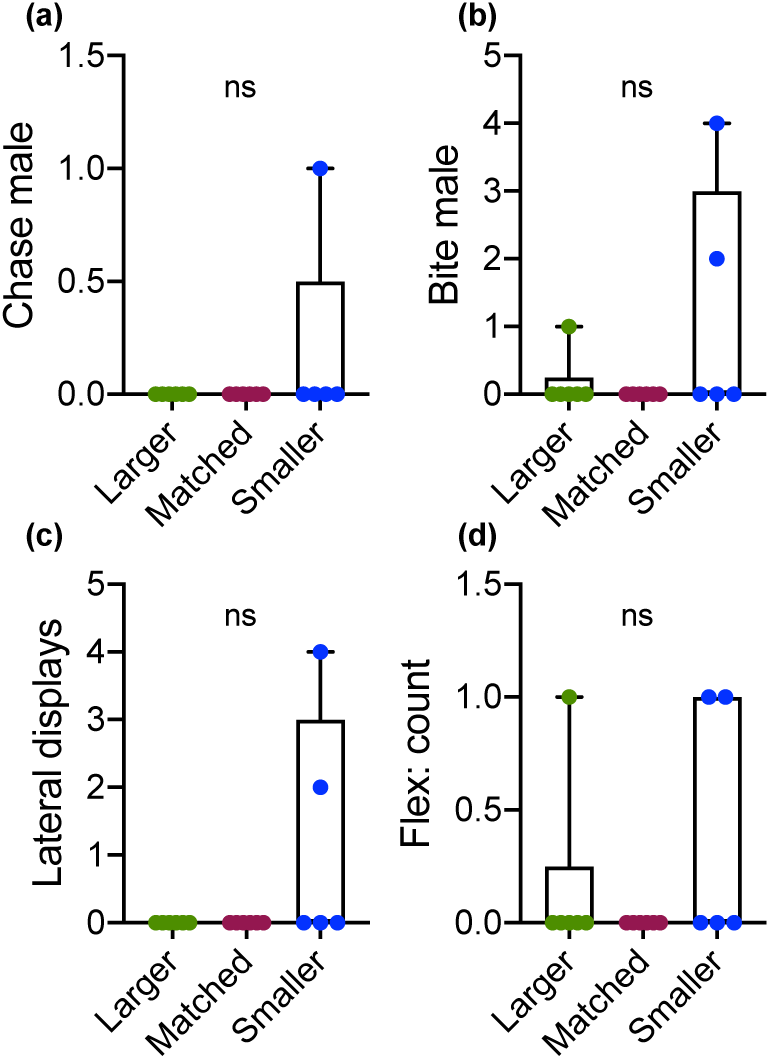
Effects of group on aggressive behavior in intruder males. (a) There were no effects of group on various intruder behaviors, including (a) chase male, (b) bite male, (c) lateral displays, and (d) flexing. Each circle represents an individual fish. Top and bottom whiskers represent maximum and minimum, respectively; top and lower boxes represent third and first quartiles, respectively; line within box represents the median.

### Correlational analyses reveal different features of resident aggression scale with relative SL

We next ran correlational analyses to assess the relationship between relative SL and behavior. These analyses showed that different aspects of aggression in the resident were altered by relative SL. For instance, the resident performed more lateral displays as relative SL increased (*r*=0.63, *N*=17, *P*=0.007) (Fig. 6a). Additionally, once relative SL reached the Matched range, residents exhibited a stark increase in the number of lateral displays they performed that continued into the larger range. The resident also flexed for longer as relative SL increased (*r*=0.53, *N*=17, *P*=0.02) (Fig. 6b). When relating the proportion of each behavior performed to the relative SL of the intruder, we found a significant positive relationship between lateral display proportion and relative SL (*r*=0.62, *N=*17, *P*=0.008) (Fig. 6c). We also found significant relationships between the latency for the resident to perform behavior the relative SL of the intruder. Specifically, residents took longer to perform the first chase at the intruder as the relative SL of the intruder increased (*r*=-0.49, *N*=17, *P*=0.04) (Fig. 6d). Residents also took longer to perform lateral displays (*r*=-0.56, *N*=17, *P*=0.02) (Fig. 6e) and flex (*r*=-0.53, *N*=17, *P*=0.02) (Fig. 6f) as the relative SL of the intruder increased.

**Figure 6.**
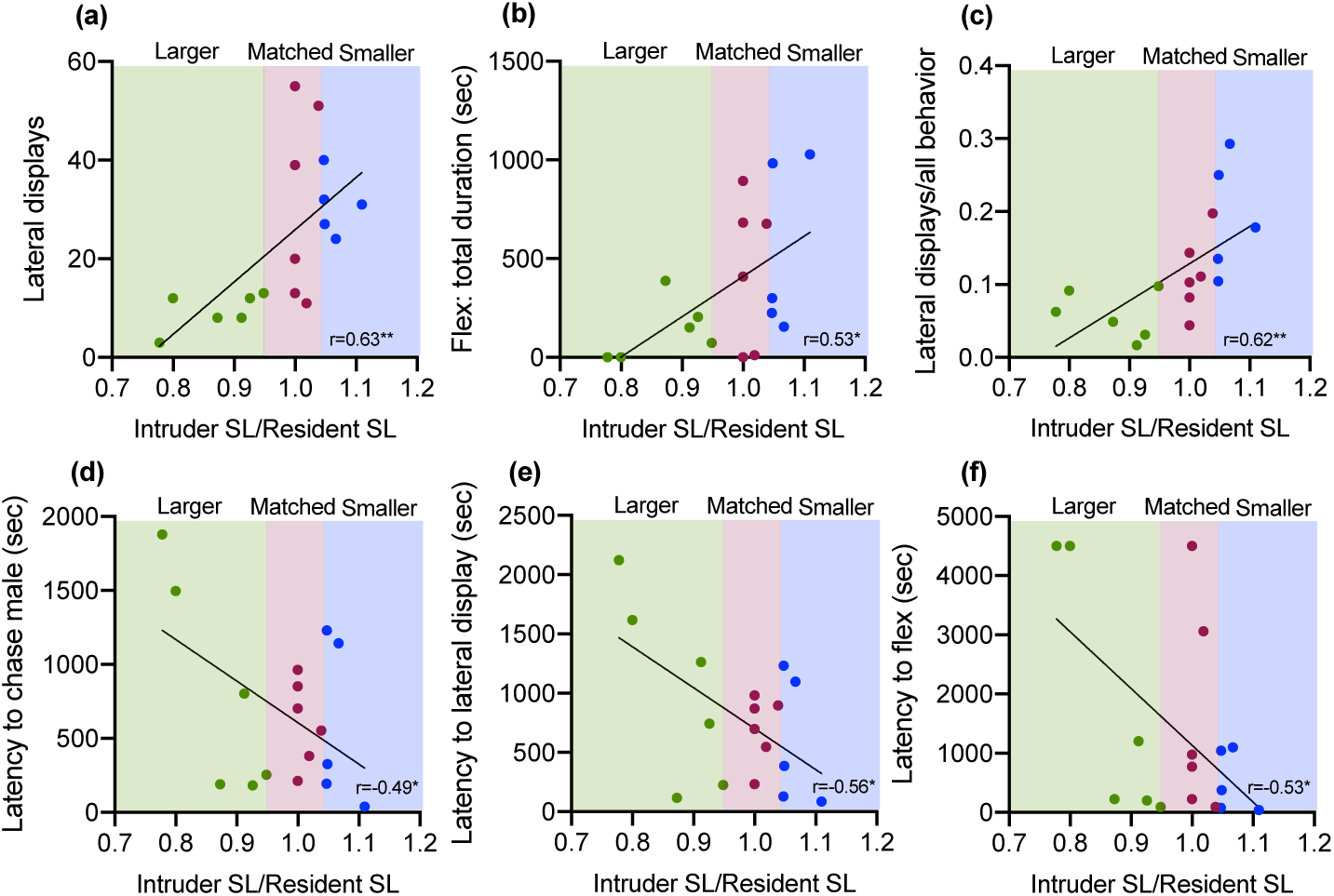
Correlations between relative standard length (SL) and behavior. Correlation analyses showed significant relationships between relative SL and several behavioral traits. (a) Larger ratios of intruder SL over resident SL were associated with more lateral displays performed by the resident. (b) Larger intruder/resident SL ratios were also associated with more time flexing by the resident male. (c) A larger proportion of behaviors performed were lateral displays when the intruder/resident SL ratio was larger. Residents took a shorter latency to (d) chase males, (e) perform lateral displays, and (f) flex when the intruder/resident SL ratio was larger. Regions of each graph shaded in green, red, or blue correspond to the different ranges of intruder/resident SL ratios on the x-axis that indicate the categorized groupings written on top of each graph (Larger, Matched, or Smaller). Pearson’s *r* values are shown in the bottom right of each correlation graph. Asterisks indicate a significant correlation. \*\**P* < 0.01; \**P* < 0.05.

### Group-level comparisons suggest residents use a different behavioral strategy depending on intruder length

We then compared resident behavior as a function of the group-level categories. There were no effects of group on physical forms of aggression (bites and chases), flexes or pot entries (*H*_*17*_ ≥4.4, *P* ≥0.12) (Fig. 7a). There was a significant effect of group on the performance of lateral displays by the resident (*H*_*17*_=8.9, *P*=0.005); both the Matched and the Smaller fish directed significantly more lateral displays at the intruder compared to the Larger fish (Fig. 7b). A significant effect of group was observed on the proportion of behaviors that were lateral displays (*H*_*17*_=9.3, *P*=0.003) (Fig. 7c). There were no effects of group on the latency to perform any specific behaviors (*H*_*17*_ ≥1.4, *P* ≥0.52).

**Figure 7.**
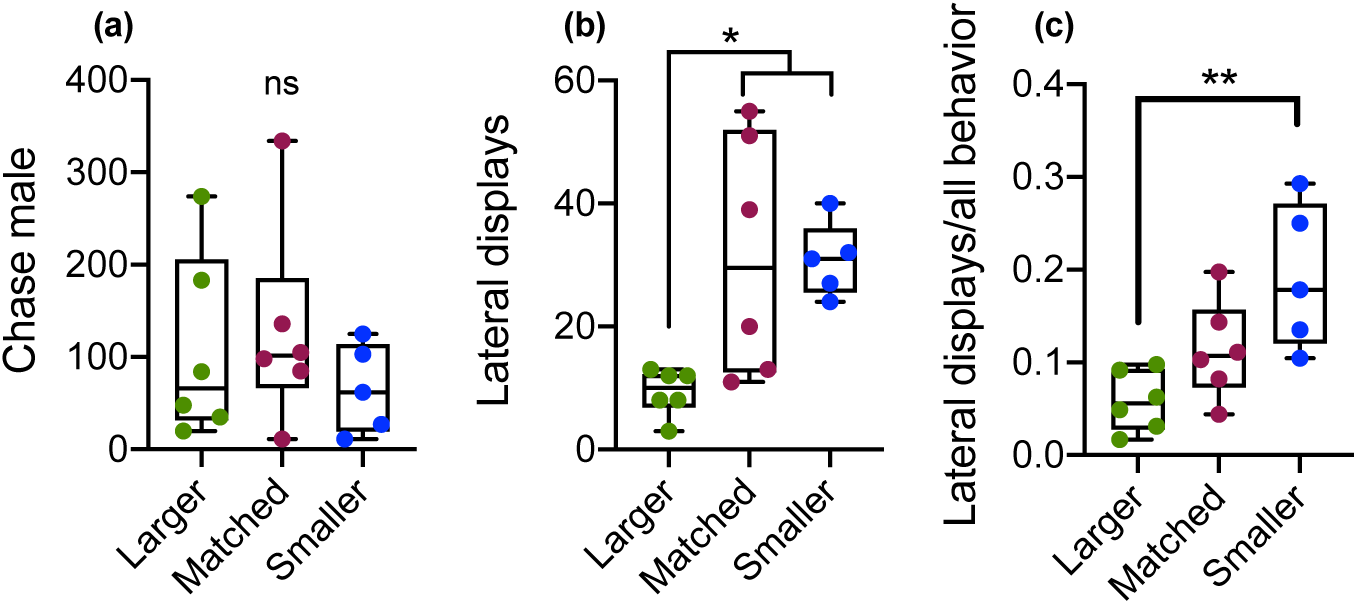
Effects of relative-size group on aggressive behavior. (a) There were no effects of group on chase male, but Matched and Smaller males performed (b) more lateral displays and (c) a larger proportion of the behaviors performed by the Smaller males were lateral displays. Each circle represents an individual fish. Top and bottom whiskers represent maximum and minimum, respectively; top and lower boxes represent third and first quartiles, respectively; line within box represents the median. \*\**P* < 0.01; \**P* < 0.05.

### No effects of gonadosomatic index or other physiological factors on resident behavior

No significant correlations were found between resident GSI and resident behaviors (*r* ≥-0.37, *N*=17, *P* ≥0.13). We also did not observe a significant effect of group on GSI (*H*_*17*_=1.3, *P*=0.53) (Fig. 8a). However, as expected, residents had significantly larger GSI than intruders (i.e., suppressed fish) (*U=*53, n_1_=18, n_2_=17, *P*=0.0006) (Fig. 8b), suggesting that residents had reached dominant social status after being placed in dominance-inducing tanks and intruders were socially suppressed.

**Figure 8.**
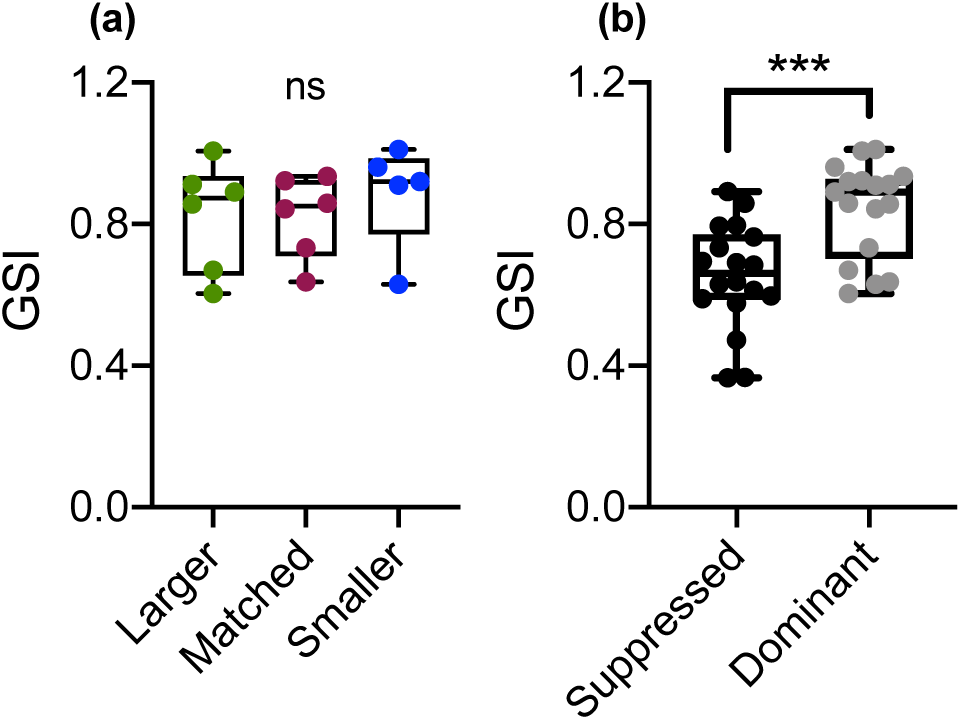
Effects of relative-size group and social status on gonadosomatic index (GSI). (a) Groups did not differ in GSI. (b) Socially suppressed males had significantly smaller GSI than dominant males. Each circle represents an individual fish. Top and bottom whiskers represent maximum and minimum, respectively; top and lower boxes represent third and first quartiles, respectively; line within box represents the median. ns=non-significant. \*\*\**P* < 0.0001.

We also assessed whether certain physiological traits in residents other than the relative SL of the intruder may have influenced our behavioral findings. Overall, while significant physiological effects were observed, they were completely unrelated to our behavioral findings. For instance, there was an effect of group on SL (*H*_*17*_=11.28, *P*=0.0003) and body mass (BM) (*H*_*17*_=11.58, *P*=0.0002), where the Smaller group had significantly larger SL and BM than the Matched group (Fig. S1a-b). Fish from the Smaller group also faced intruders that were significantly larger in terms of SL (*H*_*17*_=11.39, *P*=0.0003) and BM (*H*_*17*_=11.31, *P*=0.0003) compared to the Larger group (Fig. S1c-d). Finally, the Smaller group had larger testes than the matched group (*H*_*17*_=6.9, *P*=0.02) (Fig. S1e). This overall pattern of differences does not systematically relate to our pattern of behavioral findings (see Fig. 4-8), suggesting that our behavioral differences are specifically related to the effects of the intruder’s relative SL.

## Discussion

We have characterized a behavioral logic underlying aggression in resident dominant males in *A. burtoni*. Specifically, when resident dominant males are exposed to an intruder who is matched or larger in relative SL, they use a behavioral strategy that emphasizes non-physical aggression. On the other hand, physical aggression in resident dominant males does not vary as a function of differences in the relative SL of the intruder. Below we describe how our results contribute to our understanding of social hierarchies in *A. burtoni*.

In *A. burtoni*, size is a critical factor in determining social rank (Fernald & Maruska, 2012; Fernald, 2012). This fact is so well-established that studies aiming to include *A. burtoni* males of lower and higher ranks can reliably induce such ranks by housing fish with others that are larger or smaller for approximately two weeks or more (For examples, see Alward, Hilliard, York, & Fernald, 2019; Burmeister, Jarvis, & Fernald, 2005; Maruska, Becker, Neboori, & Fernald, 2013; Maruska & Fernald, 2010). Nevertheless, relative size in *A. burtoni* has typically been used only in this way. One reason small differences in relative size were not considered to be behaviorally relevant based on previous findings is the lack of consistency in behavioral quantification and analysis itself. For instance, different aggressive behaviors in *A. burtoni* have at times been represented by a single metric, in which both physical and non-physical aggression were treated as one measure called total aggression (for example, see Desjardins & Fernald, 2010). However, recent studies have shown that physical and non-physical aggression are uncorrelated in *A. burtoni*, suggesting that these aggressive behaviors function differently during social interactions. For instance, Loveland and colleagues showed a lack of correlation between lateral displays and border fights, a type of physical aggression (Loveland, Uy, Maruska, Carpenter, & Fernald, 2014). Additionally, a time-course study showed a robust decrease in the performance of lateral displays from morning to afternoon, without a change in the performance of border fights during the same time period (Alward et al., 2019). This finding provides further evidence that physical and non-physical aggression are dissociable in *A. burtoni*. By focusing on individual types of aggressive behavior we were able to detect fine-grained differences in behavioral output as a function of subtle differences in SL.

Our results are in line with what has been observed in other fish species. For instance, in the convict cichlid *Cichlasoma nigrofasciatum*, Iateral displays are performed less when fish could interact visually before allowed to interact physically, compared to when they could not see each other before physical interaction (Keeley & Grant, 1993). Notably, physical aggression did not differ regardless of whether the fish could see each other before being allowed to physically interact. Thus, as in *A. burtoni*, evidence exists in other fish such as *C. nigrofasciatum* that non-physical and physical aggression are used differently depending on the social environment. For *A. burtoni* specifically, lateral displays are used more frequently when SL asymmetries are smaller, suggesting lateral displays are used to settle conflicts that are difficult to resolve.

In angelfish (*Pterophyllum scalare*) larger males competing in a neutral territory always won contests (Chellappa, Yamamoto, Cacho, & Huntingford, 1999). On the other hand, when a resident-intruder asymmetry existed the resident always won irrespective of relative intruder size. Hence, as with *A. burtoni*, relative size influences behavior but residents have an advantage in resident-intruder contests. Indeed, the prior-residence advantage effect has been well demonstrated in laboratory and field situations (Alcock, 2009; Mesterton-Gibbons & Sherratt, 2016). Future studies modifying social experience of both intruders and residents in *A. burtoni* may yield novel insights into the behavioral logic of aggression in *A. burtoni*.

Our results suggest there is a complex relationship between social experience and behavioral responses to size differences in *A. burtoni*. Indeed, if it was the case that only size differences guided behavioral performance, then intruder males that were larger than the resident should have performed more lateral displays--but this was clearly not the case. These results suggest a winner and/or loser effect plays a role in guiding social decisions in male *A. burtoni*. In the winner effect, competitors who win contests are more likely to win future contests than losers (Dugatkin, 1997; Dugatkin & Earley, 2004). Here, socially suppressed males are likely to be losing contests repeatedly. Furthermore, testosterone, which is higher in dominant males than non-dominant males (Parikh, Clement, & Fernald, 2006), increases the winner effect (Oliveira, Silva, & Canário, 2009). Moreover, androgen receptor activation is required for social dominance (Alward et al., 2019). Based on the above, we hypothesize that testosterone may modulate cost thresholds in *A. burtoni* males. Future work manipulating testosterone signaling pharmacologically or genetically will be fundamental in determining the functional relationship between testosterone, winner/loser effects, and cost thresholds in *A. burtoni*.

## Conclusion

We discovered in a highly social cichlid fish that relative size differences between a dominant resident and a non-dominant intruder male affects social decisions made by the resident male. Our findings lay the foundation for future work on the different social and biological factors that may affect behavioral strategies in *A. burtoni* and add to the existing work on models of aggression in social fish species.

## Acknowledgements

We thank Christopher Skalnik for helpful feedback assisting in the experimental design and Anne Fernald and Andrew Hoadley for providing feedback on an earlier version of the manuscript.

## Supplementary Figure and Legend

**Figure S1.**
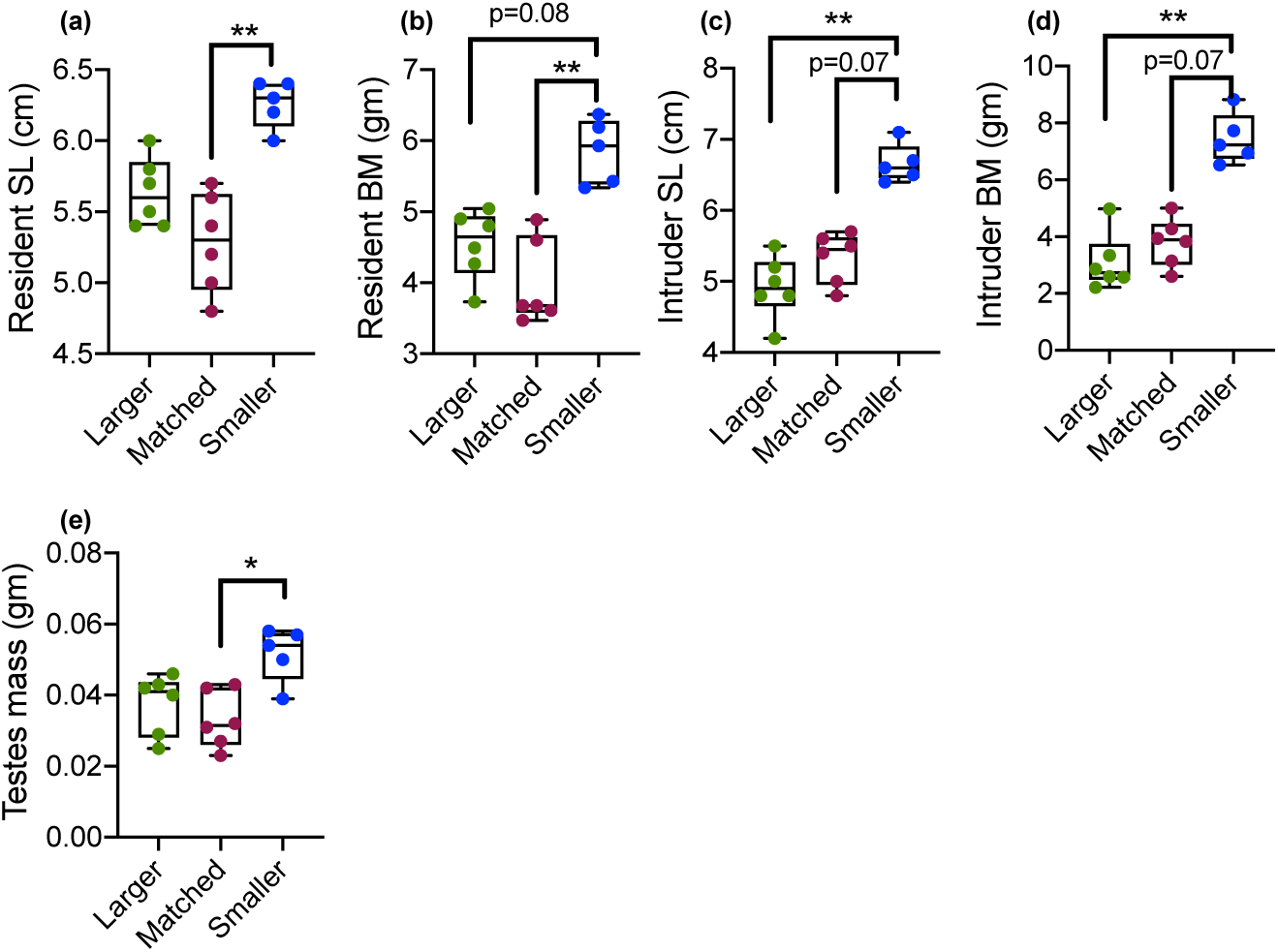
Effects of group on body size measures and testes mass. (a-e) For all measures shown the Larger and Matched group did not differ. The Smaller group males were larger than the Matched group for (a) SL and (b) BM, and (e) testes mass. (b) There was a statistical trend for the Smaller group fish to have larger BM than the Larger group fish. (c-d) Smaller group fish were exposed to larger intruders than the Larger and Matched group fish, but this was only a statistical trend for the latter group. Each circle represents an individual fish. Top and bottom whiskers represent maximum and minimum, respectively; top and lower boxes represent third and first quartiles, respectively; line within box represents the median. \*\**P* < 0.01; \**P* < 0.05.

## References

Alcazar, R. M., Hilliard, A. T., Becker, L., Bernaba, M., & Fernald, R. D. (2014). Brains over brawn: Experience overcomes a size disadvantage in fish social hierarchies. Journal of Experimental Biology, 217(9), 1462–1466. https://doi.org/10.1242/jeb.097527

Alcock, J. (2009). Animal behavior: An evolutionary approach. Sinauer Associates.

Alward, B. A., Hilliard, A. T., York, R. A., & Fernald, R. D. (2019). Hormonal regulation of social ascent and temporal patterns of behavior in an African cichlid. Hormones and Behavior, 107(December 2018), 83–95. https://doi.org/10.1016/j.yhbeh.2018.12.010

Burmeister, S. S., Jarvis, E. D., & Fernald, R. D. (2005). Rapid behavioral and genomic responses to social opportunity. PLoS ONE, 3(11), 1–9. https://doi.org/10.1371/journal.pbio.0030363

Chellappa, S., Yamamoto, M. E., Cacho, M. S. R. F., & Huntingford, F. A. (1999). Prior residence, body size and the dynamics of territorial disputes between male freshwater angelfish. Journal of Fish Biology, 55, 1163–1170.

Desjardins, J. K., & Fernald, R. D. (2010). What do fish make of mirror images? Biology Letters, 6(6), 744–747. https://doi.org/10.1098/rsbl.2010.0247

Dugatkin, L. A. (1997). Winner and loser effects and the structure of dominance hierarchies. Behavioral Ecology, 8(6), 583–587. https://doi.org/10.1093/beheco/8.6.583

Dugatkin, L. A., & Earley, R. L. (2004). Individual recognition, dominance hierarchies and winner and loser effects. Proceedings of the Royal Society B: Biological Sciences, 271(1547), 1537–1540. https://doi.org/10.1098/rspb.2004.2777

Fernald, R. D., & Maruska, K. P. (2012). Social information changes the brain. Proceedings of the National Academy of Sciences, 109(Supplement_2), 17194–17199. https://doi.org/10.1073/pnas.1202552109

Fernald, R.D. (2012). Social control of the brain. Annual Review of Neuroscience, 35, 135–151.

Fernald, Russell D., & Hirata, N. R. (1977). Field study of Haplochromis burtoni: Quantitative behavioural observations. Animal Behaviour, 25, 964–975. https://doi.org/10.1016/0003-3472(77)90048-3

Holekamp, K. E., & Strauss, E. D. (2016). Aggression and dominance: an interdisciplinary overview. Current Opinion in Behavioral Sciences, 12, 44–51. https://doi.org/10.1016/j.cobeha.2016.08.005

Keeley, E. R., & Grant, J. W. A. (1993). Visual information, resource value, and sequential assessment in convict cichlid (Cichlasoma nigrofasciatum) contests. Behavioral Ecology, 4(4), 345–349. https://doi.org/10.1093/beheco/4.4.345

Loveland, J. L., Uy, N., Maruska, K. P., Carpenter, R. E., & Fernald, R. D. (2014). Social status differences regulate the serotonergic system of a cichlid fish, Astatotilapia burtoni. Journal of Experimental Biology, 217(15), 2680–2690. https://doi.org/10.1242/jeb.100685

Maruska, K. P., Becker, L., Neboori, A., & Fernald, R. D. (2013). Social descent with territory loss causes rapid behavioral, endocrine and transcriptional changes in the brain. Journal of Experimental Biology, 216(19), 3656–3666. https://doi.org/10.1242/jeb.088617

Maruska, Karen P., & Fernald, R. D. (2010). Behavioral and physiological plasticity: Rapid changes during social ascent in an African cichlid fish. Hormones and Behavior, 58(2), 230–240. https://doi.org/10.1016/j.yhbeh.2010.03.011

Maynard Smith, J., & Harper, D. G. C. (1988). The evolution of aggression: can selection generate variability? Philosophical Transactions of the Royal Society of London. Series B, Biological Sciences, 319, 557–570.

Mesterton-Gibbons, M., & Sherratt, T. N. (2016). How residency duration affects the outcome of a territorial contest: Complementary game-theoretic models. Journal of Theoretical Biology, 394, 137–148. https://doi.org/10.1016/j.jtbi.2016.01.016

Oliveira, R. F., Silva, A., & Canário, A. V. M. (2009). Why do winners keep winning? Androgen mediation of winner but not loser effects in cichlid fish. Proceedings of the Royal Society of London B., 276(March), 2249–2256. https://doi.org/10.1098/rspb.2009.0132

Parikh, V. N., Clement, T. S., & Fernald, R. D. (2006). Androgen level and male social status in the African cichlid, Astatotilapia burtoni. Behavioural Brain Research, 166(2), 291–295. https://doi.org/10.1016/j.bbr.2005.07.011

Reddon, A. R., Voisin, M. R., Menon, N., Marsh-Rollo, S. E., Wong, M. Y. L., & Balshine, S. (2011). Rules of engagement for resource contests in a social fish. Animal Behaviour, 82(1), 93–99. https://doi.org/10.1016/j.anbehav.2011.04.003

Solomon-Lane, T. K., & Hofmann, H. A. (2019). Early-life social environment alters juvenile behavior and neuroendocrine function in a highly social cichlid fish. Hormones and Behavior, 115(April), 104552. https://doi.org/10.1016/j.yhbeh.2019.06.016

Taborsky, M. (1984). Broodcare helpers in the cichlid fish Lamprologus brichardi: Their costs and benefits. Animal Behaviour, 32, 1236–1252.

Taborsky, M. (1985). Breeder-Helper Conflict in a Cichlid Fish With Broodcare Helpers: an Experimental Analysis. Behaviour, 95, 45–75.

van Staaden, M. J., Searcy, W. A., & Hanlon, R. T. (2011). Signaling aggression. In Advances in Genetics (1st ed., Vol. 75). https://doi.org/10.1016/B978-0-12-380858-5.00008-3

